# A Markov Random Field Model for Network-based Differential Expression Analysis of Single-cell RNA-seq Data

**DOI:** 10.1101/2020.11.11.378976

**Authors:** Hongyu Li, Biqing Zhu, Zhichao Xu, Taylor Adams, Naftali Kaminski, Hongyu Zhao

## Abstract

**Background:** Recent development of single cell sequencing technologies has made it possible to identify genes with different expression (DE) levels at the cell type level between different groups of samples. In this article, we propose to borrow information through known biological networks to increase statistical power to identify differentially expressed genes (DEGs).

**Results:** We develop MRFscRNAseq, which is based on a Markov Random Field (MRF) model to appropriately accommodate gene network information as well as dependencies among cell types to identify cell-type specific DEGs. We implement an Expectation-Maximization (EM) algorithm with mean field-like approximation to estimate model parameters and a Gibbs sampler to infer DE status. Simulation study shows that our method has better power to detect cell-type specific DEGs than conventional methods while appropriately controlling type I error rate. The usefulness of our method is demonstrated through its application to study the pathogenesis and biological processes of idiopathic pulmonary fibrosis (IPF) using a single-cell RNA-sequencing (scRNA-seq) data set, which contains 18,150 protein-coding genes across 38 cell types on lung tissues from 32 IPF patients and 28 normal controls.

**Conclusions:** The proposed MRF model is implemented in the R package MRFscRNAseq available on GitHub. By utilizing gene-gene and cell-cell networks, our method increases statistical power to detect differentially expressed genes from scRNA-seq data.

## Background

With recent advancement in single-cell RNA sequencing (scRNA-seq) technologies, it has opened up unique opportunities to understand genomic and proteomic changes at the single cell resolution. Such data allow us to identify cell type specific differentially expressed genes (DEGs) that are associated with diseases, e.g. idiopathic pulmonary fibrosis (IPF). There exist many statistical methods that can perform DE analysis for scRNA-seq data. First of all, traditional statistical methods such as two-sample *t*-test, Wilcoxon rank sum test, logistic regression, and negative binomial regression are widely used to detect DEGs for scRNA-seq data. DE methods that are tailored for scRNA-seq data have also been developed over the last decade. For instance, Single-Cell Differential Expression (`SCDE`) [1] fits a mixture of Poisson model and Negative Binominal model for the zeros and positive mean expressions separately. Model-based Analysis of Single-cell Transcriptomics (`MAST`) [2] utilizes a two-part hurdle model to simultaneously model the rate of expression and the mean expression level. One novel aspect of `MAST` is that it adjusts the fraction of genes expressed across cells to obtain more reliable estimates. `scDD` [3] uses a conjugate Dirichlet process mixture to identify DEGs. `DESingle` [4] adopts a zero-inflated negative binomial distribution to model count data and identify DEGs. Meanwhile, nonparametric methods such as `SigEMD` [5], `EMDomics` [6], and `D3E` [7] have also been developed for scRNA-seq data to detect DEGs. Review papers [8, 9] have pointed out that methods that were tailored for scRNA-seq data do not show significantly better performance compared to traditional methods. Surprisingly, traditional methods such as *t*-test and Wilcoxon test also have fairly robust performance.

In addition, several papers [10, 11, 12] have pointed out that variations between biological replicates should be properly controlled when performing DE analysis, i.e., individual effects. As a result, the family of pseudo-bulk methods are also commonly used in the DE analysis. These methods usually aggregate cell-level counts into sample-level pseudo-bulk counts, and then use methods that were originally proposed for bulk RNA-seq data to detect DEGs, such as `edgeR` [13], `DESeq2` [14], and `limma` [15]. On the other hand, methods based on mixed models [11] have also been proposed to capture the random effects for individuals. However, Crowell [10] showed in their comparison analysis that the mixed model-based methods did not perform significantly better compared to the aggregation-based pseudo-bulk methods. Moreover, these mixed model-based methods also require larger computational resources and longer computational time, which may not be worthwhile. Currently the detection of DEGs for scRNA-seq data still remains a challenge. Nonetheless, although it is well known that genes and cells do not work independently, none of the existing methods take gene network information or dependencies among cells into consideration. Given the large scale and complexity of the scRNA-seq data, one key challenge is how to appropriately accommodate these dependencies to better identify cell-type specific DEGs.

In this paper, we propose a Markov Random Field (MRF) model that can capture gene network and cell type information. We note that the MRF model has been applied to both genome-wide association studies and bulk RNA-seq studies to model dependencies in genomic and transcriptomic data. For instance, biological pathways were used to model the structure of neighboring genes [16, 17, 18]. In addition, the similarities between brain regions and adjacent time points were incorporated to jointly model the spatial-temporal dependencies for human neurodevelopment data [19]. Our method adopts local false discovery rate framework that was developed by Efron [20] to identify cell-type specific DEGs. We implement an efficient EM algorithm [21] with mean field-like approximation [22, 23, 24] to estimate model parameters. Then we utilize Gibbs sampler to estimate the posterior probabilities to infer cell-type specific DE status.

We applied our method to a recent study that collected scRNA-seq data using lung tissues from 32 IPF patients and 28 normal controls [25]. The objective of the analysis is to detect cell-type specific DEGs between IPF patients and normal controls. Idiopathic pulmonary fibrosis (IPF) is an incurable aggressive lung disease. It progressively scars the lung and causes usual interstitial pneumonia (UIP).

However, to date, what causes the scarring remains unknown. IPF affects around three million people globally [26, 27], with its mortality rate much higher than many cancers, and the median survival time for patients without a lung transplant is about three to four years [28, 29]. Many efforts have been made to understand the pathogenesis and biological processes of this disease. For instance, genome-wide association studies (GWAS) have identified 20 regions in the human genome that are associated with increased risk to IPF [30, 31, 32, 33]. In addition, transcriptome analyses through microarrays [34, 35, 36, 37] and RNA-seq [38, 39, 40] have revealed genes and pathways related to IPF. In particular, a recent review described in detail how transcriptome analyses helped to identify novel genes involved in the pathogenesis of IPF and the importance of using single-cell RNA-seq analysis to discover cell-type specific DEGs [41]. In order to assess the performance of our proposed MRF model, simulation study was conducted under various scenarios. The results for simulation study and the DE results for the IPF scRNA-seq analysis are shown in the third section. We conclude the manuscript with a brief discussion in the last section.

## Methods

### Markov Random Field Model

#### Model Setup

Given normalized single cell RNA-seq data, let *y_gcpk_* denote the normalized observed expression of gene *g* in cell type c in the *k*^th^ replicate in condition *p*. We let *G* denote the number of genes and *C* denote the number of cell types. For simplicity, we assume *P* = 2. We assume that the cells have been correctly assigned to their corresponding cell types. In each group, there are *n_gcp_* samples for the *p*^th^ group (either disease or control group). The number of samples here is the number of cells belonging to this cell type. Let **y**_*gc*1_ and **y**_*gc*2_ denote the vectors of expression values for gene *g* in cell type *c* for the two groups. The two-sample *t*-statistic can be constructed as

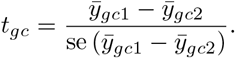

Then we transform the test statistic into z-scores,

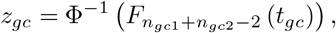

where *n*_*gc*1_ and *n*_*gc*2_ are the number of samples for the two groups, e.g. disease and control groups, for gene *g* in cell type *c*; Φ is the cumulative distribution function of a standard normal distribution; and *F* is the cumulative distribution function for a student-*t* distribution with *n*_*gc*1_ + *n*_*gc*2_ − 2 degrees of freedom. The gene expression data are then represented by a summary statistic matrix **Z**, where each entry *z_gc_* represents the evidence of differential expression between the two groups for each gene across cell types. **Z** is a *G* × *C* matrix. Let *w_gc_* denote the binary latent state representing whether gene *g* is differentially expressed in cell type *c* between the two groups. Then **W** is the latent state matrix, which has the same dimension as **Z**. Because *w_gc_* has two states, we assume that *z_gc_* follows a mixture distribution,

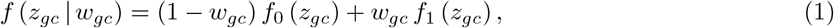

where *f*_0_(*z_gc_*) is the null density and *f*_1_(*z_gc_*) is the non-null density. The null and non-null densities are estimated through Efron’s nonparametric empirical Bayes framework [20]. The inference on the latent state **W** is our primary objective. In the following, we construct the MRF model that accommodates cell type dependencies and gene network information. A gene network information is represented by an undirected graph, with a set of nodes 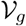, which correspond to cell-type specific genes, and a set of edges 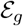, which represent the relationships among the nodes. For each gene *g*, we can use the following vector to denote its cell-type specific DE status,

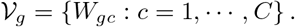

The set of edges 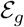 can be divided into two subsets, 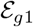 and 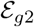. For two genes *g* and *g*′, if there is a known relationship, e.g. from a pathway database, we write *g ~ g*′. For a given gene *g*, let 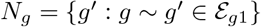 be the set of genes that have known relationships with this gene. Similarly, for two cell types *c* and *c*′, if there is a known relationship, we write *c ~ c*′. For a given cell type *c*, let 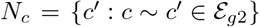 be the set of cell types that have close relationships with cell type *c*. Then we can write two sets of edges as

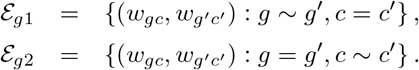

Therefore, edges in 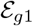 capture similarities between genes based on gene network information, while edges in 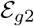 capture the dependencies between cell types. Then we construct a pairwise interaction MRF model [42],

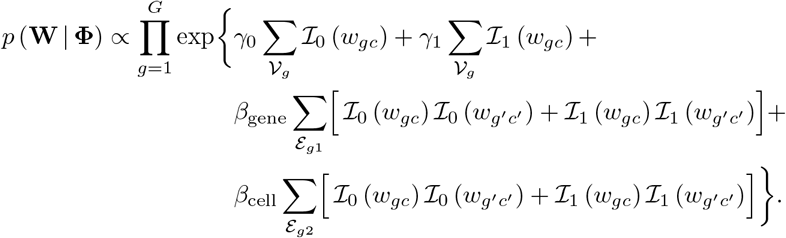

Here 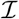 is an indicator function, i.e., when *w_gc_* = 1, 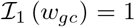. Let *γ* = *γ*_1_ – *γ*_0_, the conditional probability for the cell-type specific DE status is

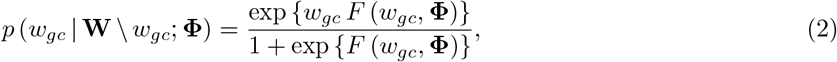

where

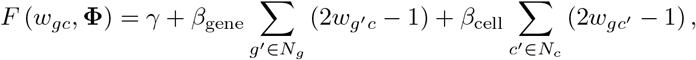

where “\” denotes other than; **Φ** = (*γ, β*_gene_, *β*_cell_). Here *β*_gene_ is the parameter that captures the similarities between genes, and *β*_cell_ is the parameter that captures cell type dependencies.

#### Parameter Estimation

For parameter estimation, we adopt the EM algorithm [21] with mean field-like approximation [22, 23, 24]. Let 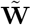 denote a configuration, the joint distribution *p*(**W** | **Φ**) can be estimated by

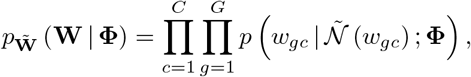

where 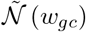 represents the neighbors, the nodes that are directly connected to this gene *g* in cell type *c*, of *w_gc_* corresponding to the fixed configuration 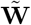. The complete log likelihood is

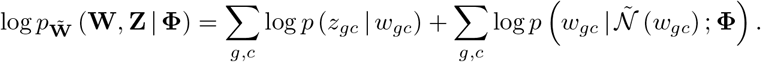

The Q function in the EM algorithm [21] is

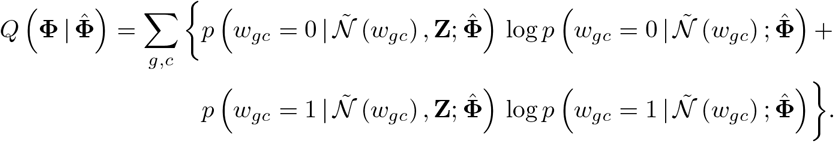

Then we use the following EM algorithm to estimate the model parameters

1. Estimate *f*_0_ and *f*_1_ using R package `locfdr` based on the z-scores. Then obtain 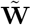 using the mixture model by Equation (1);
2. Expectation-step: Let 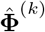 be the estimated parameters in the *k*^th^ iteration. The *Q* function 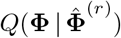 can be calculated from the fixed configuration 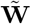;
3. Maximization-step: Update **Φ** with 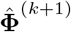, which maximizes 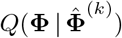;
4. Obtain the updated 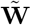 by sequentially updating 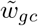 by the Gibbs sampler with posterior probability proportional to

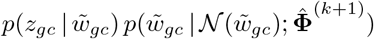
5. Repeat steps 2-4 until convergence.

#### Detecting Differentially Expressed Genes

After we obtain the model parameters from the EM algorithm, we use a Gibbs sampler to sample the posterior probabilities. We then use the posterior probability-based definition of FDR [43, 44] to identify DEGs. We first estimate the posterior local FDR *p_gc_* using Gibbs sampler,

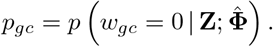

Then we sort *p_gc_* in ascending order, and let *p*_(*i*)_ denote the sorted values. We find *k* such that

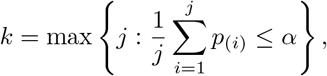

and we reject the first *k* null hypotheses.

### IPF scRNA-seq Data Analysis

The gene expression levels were profiled in each single cell in the IPF scRNA-seq dataset. We used R package `Seurat` [45] to perform data preprocessing and quality control. Specifically, cells that had unique gene counts greater than 5,000 or less than 200 were filtered out. Cells that had more than 5% mitochondrial counts were also excluded from further analysis. After quality control, there were 18,150 protein-coding genes profiled in 114, 364 cells. We normalized the expression data for each cell by the total expression multiplied by a scale factor of 10,000 and then log-transformed the results. Each single cell corresponds to a cell type. Since about 87% of the cells were myeloid and lymphoid cells, we focused on the immune cells in further analyses. We used UMAP [46], which is a manifold learning technique for dimension reduction, to plot the cell types. There were 18 distinct immune cell types (Figure 1A).

**Figure 1.**
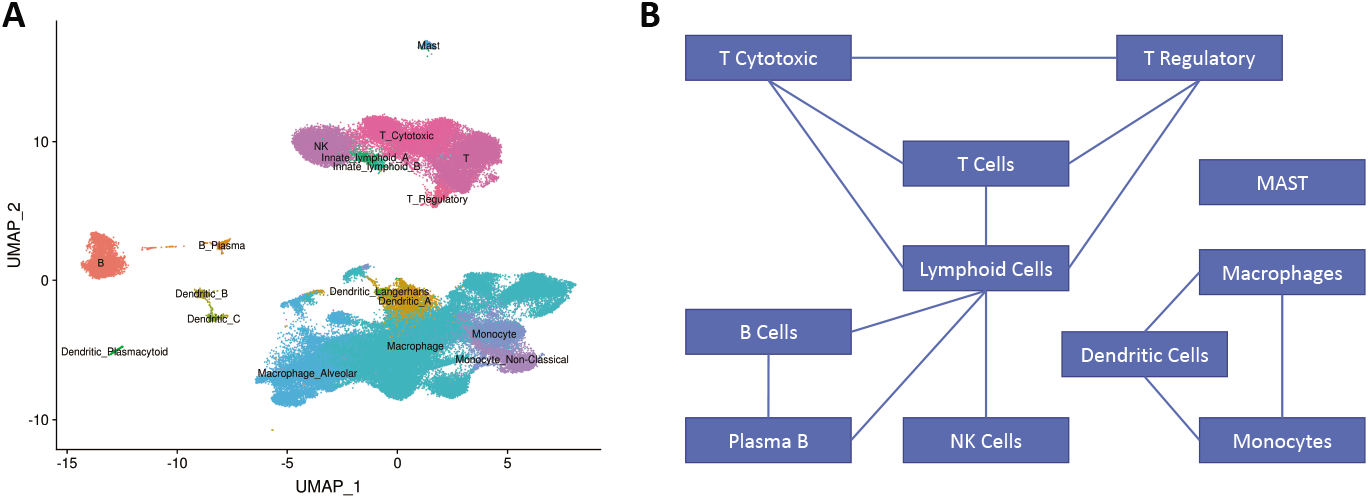
Cell Type Clusters. **(A)** shows the UMAP of eighteen immune cell types, and **(B)** shows the network among these immune cell types that are determined by domain knowledge.

In this paper, instead of considering a network with 18,150 genes across 18 cell types, we focused on 2,000 genes that exhibited high cell-to-cell variation between cell types. Previous research [45, 47] showed that focusing on these highly variable genes in DE analysis helps to highlight significant biological signals. We extracted the gene network information from two well-known protein-protein interaction network (PPIN) databases, BioGrid [48] and IntAct [49]. For these 2,000 highly variable genes, the BioGrid database have 5,400 edges, while there are 3,104 edges in the IntAct database. The two gene networks are visualized in Supplementary Figure 1 [see Additional file 1]. There is an overlap of 1,754 edges between the two databases. In addition, the dependencies among cells were determined by domain knowledge (Figure 1B).

For these 2,000 genes across 18 cell types, we fitted two separate MRF models utilizing gene networks from BioGrid and IntAct, respectively. We obtained the parameter estimates by implementing the EM algorithm with 200 iterations. With parameter estimates fixed, we then ran 20,000 iterations of Gibbs sampler with 10,000 iterations as burn-in to obtain posterior probabilities. These were the two main models of our analysis, and we labeled them as Main model with BioGrid and Main model with IntAct, respectively. In addition, the model that used only two-sample *t*-test statistics (no gene and cell networks) was labeled as Main model, which was not an MRF model.

In addition, we considered two sets of supplementary models. First, in order to assess the importance of cell network structures in the MRF model, we considered two additional cell networks C1 and C2 (Supplementary Figure 3) [see Additional file 1]. We used the same observed DE evidence as well as the gene network structures in the Main models, and fitted four additional MRF models with cell networks C1 and C2. We labeled these models with the additional annotation of C1 and C2, respectively. Second, our method can be generalized to use observed DE evidence from other existing DE methods in addition to the two-sample *t*-test. Most DE methods for scRNA-seq data typically output *p*-values and log-fold changes for their differential expression analysis. Our model can be readily applied to these existing models with ease. We chose the following two methods as examples: Wilcoxon and MAST. Based on the simulation results, these two methods generally have stable and robust performance, especially when the number of cells in each subject is small. In our IPF scRNA-seq data, many cell types have a limited number of cells per subject, i.e., T Regulatory cells and B Plasma cells, so these two methods are more suitable in our IPF analysis. We used `Seurat` to obtain DE results for these two models. Then we transformed the *p*-values to *z*-scores and used the sign of log-fold changes to determine the sign of *z*-scores. With an additional set of observed DE evidence (*z*-scores), we used the same gene sets and biological networks in the Main models to fit four additional MRF models. We labeled these models accordingly. For instance, the model that only considered DE results from `MAST` analysis was labeled as `MAST` model, and the MRF model that incorporated `MAST` DE results with BioGrid gene network was labeled as `MAST` with BioGrid. A table summarizing the models we used in this manuscript is provided in Supplementary Table 1 [see Additional File 1].

### Simulation Study

Simulation study was conducted to assess the performance of our proposed MRF procedure. The single cell RNA-seq data are count data with zero inflation due to dropout and over-dispersion; thus, a zero-inflated negative binomial (ZINB) distribution is suitable to model the read counts and excessive of zeros in the scRNA-seq data. The zero-inflated negative binomial distribution consists of three parameters: mean, dispersion, and inflation. We obtained these parameters based on the IPF scRNA-seq data. In details, for each cell type, we fitted a ZINB distribution across 2, 000 highly variable genes. We obtained three vectors of estimated parameters for the mean, dispersion, and inflation of length 2, 000. Then we fitted a gamma distribution for the estimated means, a gamma distribution for the estimated dispersion parameters, and a beta distribution for the estimated inflation parameters.

We repeated this for eighteen cell types; thus, for each cell type, we had an estimated gamma distribution for the mean, an estimated gamma distribution for the dispersion, and an estimated beta distribution for the inflation.

For our simulated data, the number of genes was set at 1, 000, and we set the number of cell types to be 18, which was the same as that in the IPF scRNA-seq data. Two groups were considered. We set the number of subjects in each group to be 15, and the number of cells for each cell type in each subject to be 50. We also varied the number of subjects and number of cells in the subsettings. The dimension of our simulated data thus was `# genes × # cell types × # groups×# subjects per group×# cells per cell type per subject`. For network structures, the cell network was set to be the same as that in the IPF scRNA-seq analysis (Main models). For gene network, we randomly selected *η* percentage genes to be connected. We varied *η* = 0.2,0.4,0.6,0.8 to reflect different proportions of connections in the gene network. For differential expression, we first randomly selected *κ* percentage genes to be DE, which gave us the initial states. With the initial states, we then used the gene and cell network structures to get the latent states by a Gibbs sampler. In each round of Gibbs sampling, the latent states were updated according to Equation (2). We varied *κ* = 0.2, 0.4 to reflect different proportions of DEGs in each setting. Then we simulated the expression count data with the zero-inflated negative binomial distribution with mean *μ_c_*, dispersion *ϕ_c_*, and inflation *γ_c_*. The three parameters were sampled from the fitted gamma and beta distributions as mentioned before, and they were cell-type specific and subject-specific. For each individual in the control group, the expression data for each gene were generated from ZINB (*μ_c_, ϕ_c_, γ_c_*). For the case group, if the latent state was 0, the count data were generated from ZINB (*μ_c_, ϕ_c_, γ_c_*); if the latent state was 1, the count data were generated from ZINB (*τ · μ_c_, ϕ_c_, γ_c_*), where *τ* = λ and 1/λ with equal probability. We chose λ to be 2. We performed the same preprocessing and quality control procedures as with the IPF single cell analysis. Then we used test statistics from two-sample *t*-test as the input of observed DE evidence, and incorporated the above simulated gene/cell networks to construct the MRF model. In order to reflect the role of the number of subjects or cells in our simulations, we considered two sub-scenarios by varying the number of subjects in each group (case/control) to be 30 (Scenario A) and the number of cells per cell type in each subject to be 100 (Scenario B). In addition, we also considered the case when λ is 3 (Scenario C). For each setting, we repeated the simulation 100 times.

We compared the results from our proposed MRF model with nine other methods: the two-sample *t*-test; the Wilcoxon Rank Sum test; the likelihood ratio test that adopts a logistic regression framework (LR); `MAST`; three pseudo-bulk methods: `DESeq2, edgeR`, and `limma`; and two mixed model methods, `dream` and its updated version `dream2. dream` and `dream2` [50] were originally designed for bulk RNA-seq studies with repeated measurements and were used in the comparative analysis by Crowell et al. [10]. The original version of `dream` uses `voom’s` [51] precision weights. In its updated version, `dream2`, it adopts the new weighting scheme in the `variancePartition` [52]. We chose these methods because they are representative and have shown fairly robust performance as noted in the review papers [8, 9, 10]. We believe that this was sufficient to demonstrate the performance of our method.

## Results

### Simulation Study

The simulation results are shown in Figure 2, and the results of the three subsettings (Scenarios A, B and C) are shown in Supplementary Figure 2A-C [see Additional file 1]. The adjusted *p*-value was set at 0.05. Sensitivity is the fraction of DEGs correctly identified by the method; specificity is the fraction of EE genes identified correctly; and FDR is calculated by the ratio of number of false positives to the number of DEGs identified. Each box-plot represents 100 replications. We note that our proposed MRF model performed significantly better than the other nine methods in terms of sensitivity. The traditional methods have fairly robust performance. The three pseudo-bulk methods have slightly lower specificity and higher FDR compared with other methods. From the two sub-settings, we note that when the number of subjects or number of cells per cell type increases, the performance of the pseudo-bulk methods and methods based on mixed models also increases as expected. Our proposed MRF model still outperforms under these sub-settings. Methods like `MAST` and Wilcoxon still have fairly robust performance, especially when the number of cells in each subject is small. In addition, in order to assess the impact of different thresholds of adjusted *p*-values on sensitivity, specificity, and FDR, and to see the trade-off directly, we chose three other adjusted *p*-value cutoffs: 0.01, 0.1, and 0.2 in addition to 0.05. In Supplementary Figure 2D [see Additional file 1], we plot sensitivity, specificity, and FDR for our proposed MRF model for the eight cases (corresponding to Figure 2) under different adjusted *p*-value cutoffs.

**Figure 2.**
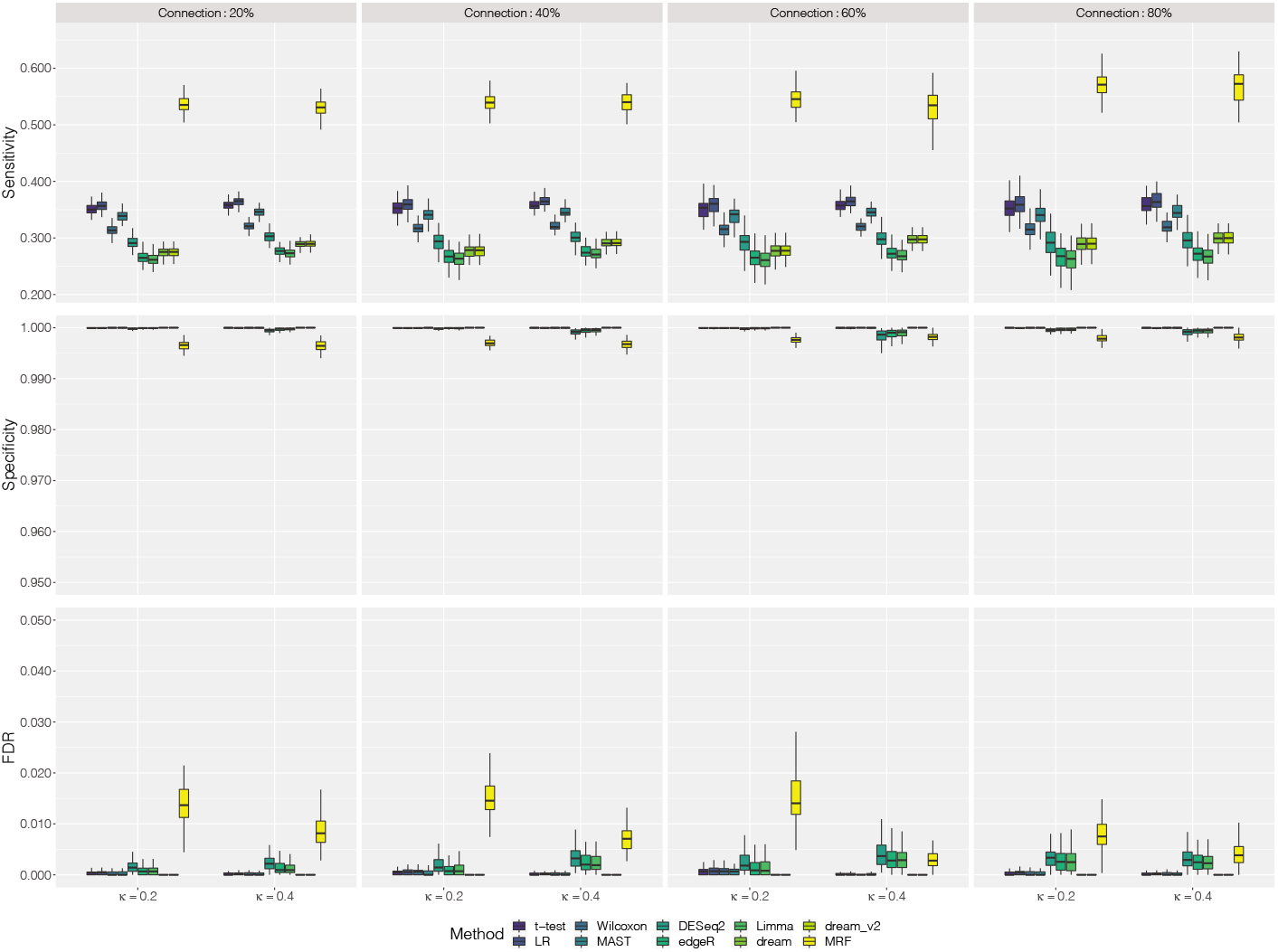
Simulation Results. The results are plotted in terms of sensitivity, specificity and FDR for two-sample *t*-test, the likelihood ratio test that adopts a logistic regression framework, Wilcoxon rank sum test, `MAST`, three pseudo-bulk methods: `DESeq2, edgeR`, and `limma`, two mixed model methods: `dream` and its updated version `dream2`, and the proposed MRF model. Each box-plot represents 100 replications.

We note that when the adjusted *p*-value threshold increases, sensitivity increases and specificity decreases for all eight cases as expected. Our proposed MRF model achieved the desired FDR control.

### IPF scRNA-seq Data Analysis

For the Main model with the BioGrid gene network, the estimated parameters were *γ* = −0.33, *β*_gene_ = 0.18, and *β*_cell_ = 0.22, whereas for the Main model with the IntAct gene network, the estimated model parameters were *γ* = −0.38, *β*_gene_ = 0.26, and *β*_cell_ = 0.22. We note that *β*_gene_ and *β*_cell_ were comparable here in both models. For DE analysis, we set the significance level at *α* = 0.01 and the corresponding posterior probability cutoff was around 0.91 for both models. Out of 2,000 genes across 18 cell types, the Main MRF model with BioGrid gene network identified 1,605 genes that were found DE in at least one cell type. For the IntAct gene network, the Main MRF model identified 1,601 genes that were DE in at least one cell type. We compared these results with two-sample *t*-tests using the Benjamini and Hochberg’s procedure for FDR, which identified 1,472 DEGs. In addition to *t*-test statistics as input for observed DE evidence, we listed the number of DEGs identified by the Wilcoxon test, and MAST, and their corresponding MRF models in Table 1. The parameter estimates for these additional models were shown in Supplementary Figure 4 [see Additional file 1]. We compared cell-type specific DEGs inferred by the model using the original test statistics alone, and two MRF models with the BioGrid and IntAct databases for three types of test statistic inputs: the Student’s *t*-test, the Wilcoxon rank sum test, and MAST in Figure 3A. We also compared DEGs identified across three types of test statistics. The UpSet [53] plot (Figure 3B) shows the overlap of DEGs identified across different models. We discovered that a major proportion of genes were found DE in all models. The detailed gene lists are provided in the Supplementary Excel file [see Additional file 3].

**Figure 3.**
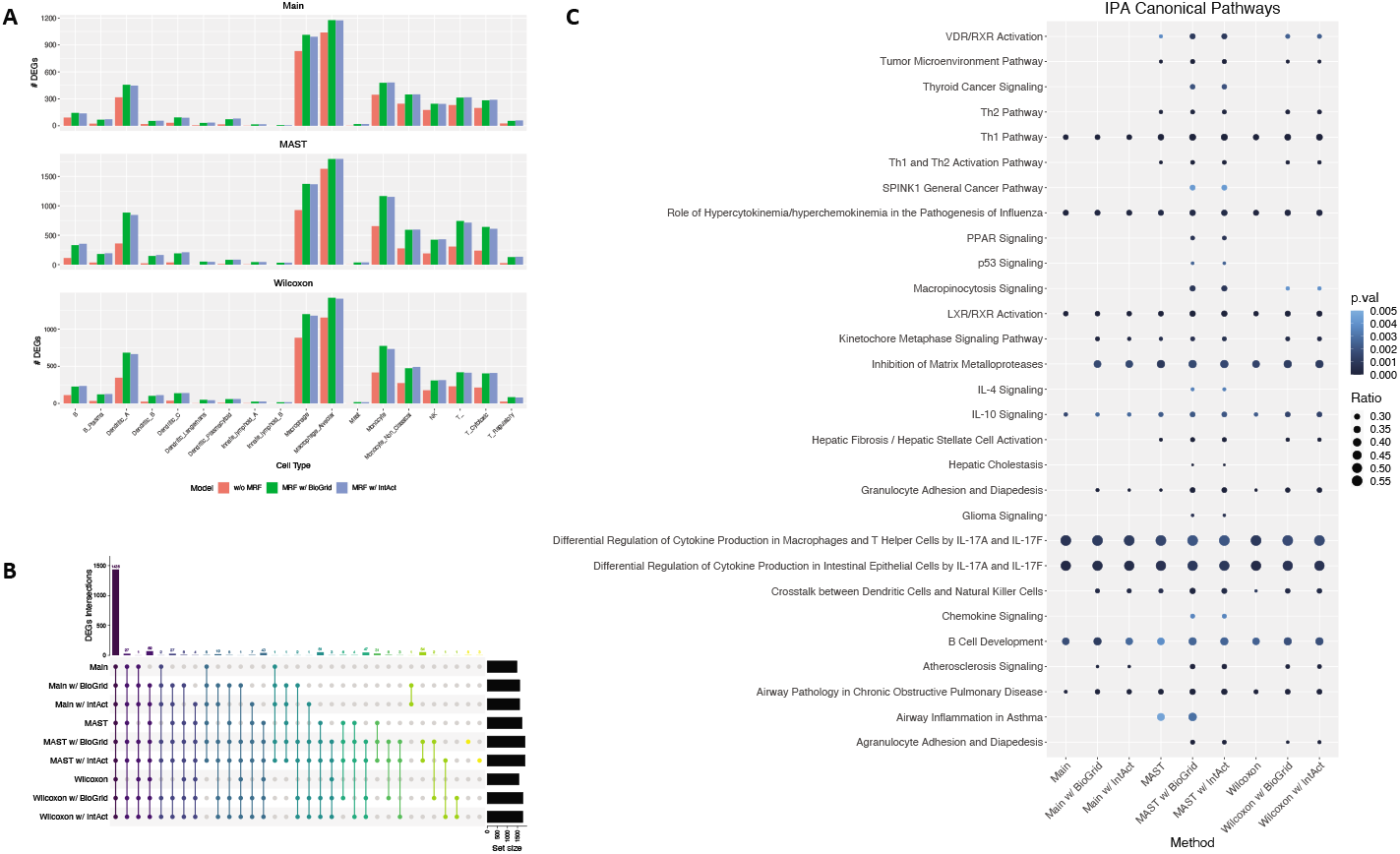
Results from IPF scRNA-seq Data Analysis. **(A)** Number of cell-type specific DEGs identified using original test statistics alone, and two MRF models with the BioGrid and IntAct database for four types of test statistic inputs: two-sample *t*-test, the Wilcoxon rank sum test, and MAST. For instance, the upper-left figure in **(A)** shows the number of cell-type specific DEGs inferred using two-sample *t*-test alone (w/o MRF), and the DE results from two MRF models that incorporated *t*-test statistics as observed DE evidence and two gene network structures (MRF w/ BioGrid and MRF w/ IntAct, respectively). **(B)** The UpSet plot shows the overlap of DEGs identified by the original models and our proposed MRF models utilizing the BioGrid and IntAct gene networks. **(C)** The top canonical pathways from Ingenuity Pathways Analysis. The *p*-value cutoff was set at 0.005 and gene ratio cutoff was set at 0.25 in order to better visualize top enriched pathways.

**Table 1.**
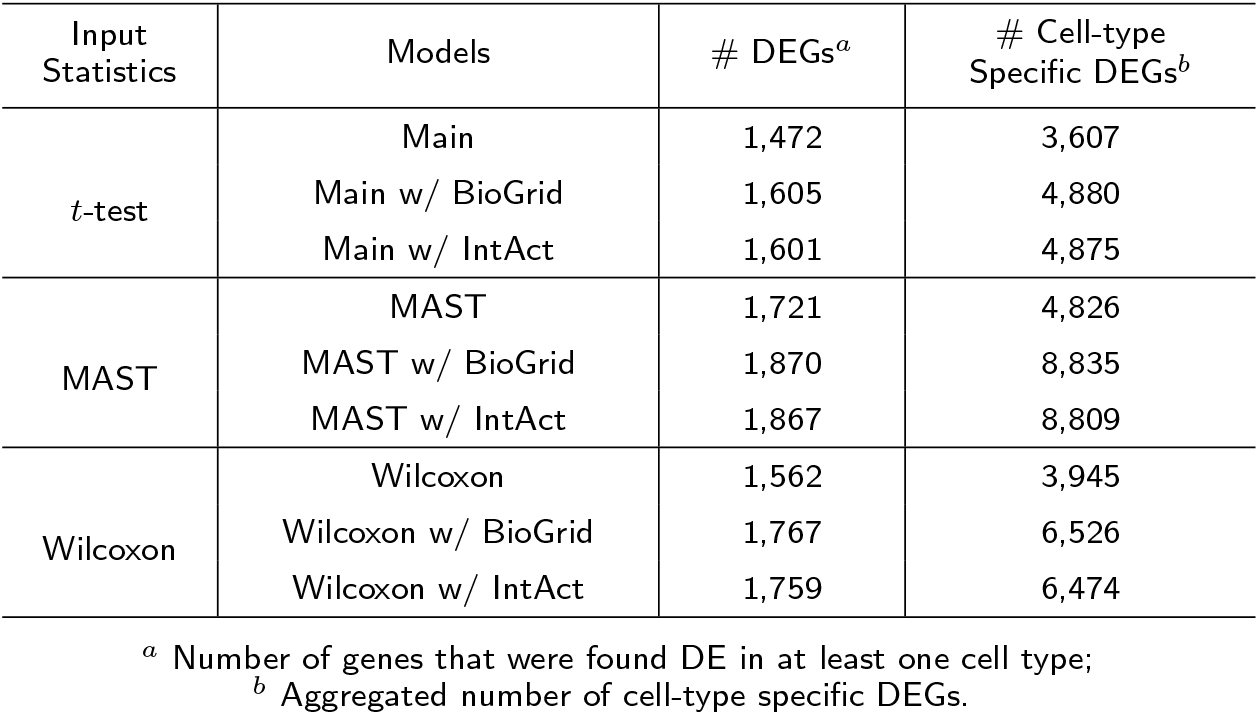
Number of DEGs Identified Under Three Model Settings with Three Types of Test Statistics

For different test statistics as observed DE evidence, our model was able to identify an additional set of novel DEGs utilizing gene-gene and cell-cell networks. We performed pathway analysis using Ingenuity Pathways Analysis (IPA, QIAGEN Inc.) [54] to get better biological insights with the inferred DEGs. In order to better visualize top enriched pathways, the *p*-value cutoff was set at 0.005 and gene ratio cutoff was set at 0.25. The complete list of pathways enriched by each model is provided in Supplementary Excel File [see Additional file 4]. Based on the IPA results in Figure 3C, we found that most pathways were enriched in all twelve models, and our proposed MRF models were able to identify an additional set of pathways that were related to IPF. In fact, we saw that a number of pathways that were related to the T helper cells were found significantly associated with IPF, which was not a surprise based on previous research [55, 56, 57, 58, 59]. In particular, three pathways: Th1 Pathway, Differential Regulation of Cytokine Production in Macrophages and T Helper Cells by IL-17A and IL-17F pathway, and Differential Regulation of Cytokine Production in Intestinal Epithelial Cells by IL-17A and IL-17F were found to be significantly associated with IPF under all models. Previous studies [55, 60, 61] showed that IL-17, the cytokines produced by the Th17 cells, participated in the development and progression of pulmonary fibrosis diseases. In addition, our proposed MRF model was able to identify the additional Th2 Pathway, and Th1 and Th2 Activation Pathway to be statistically significant. Previous findings [58, 62] showed that the Th2 cells stimulated fibroblast proliferation and activation, and promoted pulmonary fibrosis. In fact, Th2 responses were related to a number of pulmonary diseases. In addition, the VDR/RXR Activation pathway was also found significant in our MRF models, which was previously found significantly associated with IPF by Boon et al [63]. The authors also noted that in a mouse study [64], the VDR-deficient mice failed to develop experimental allergic asthma, and this suggested that vitamin D play a key role in the generation of Th2-driven inflammation in lung diseases. Moreover, the MRF model also identified the hepatic fibrosis and hepatic stellate cell activation pathway. Tsuchida et al. [65] noted in their paper that this pathway was well-established as the central driver of hepatic fibrosis in human liver. In addition, the authors discovered that HSC-specific deletion of inte-grin *αυ* protects mice from hepatic, renal and pulmonary fibrosis. To sum up, the inferred canonical pathways from our approach have biological meanings and are strongly related to IPF.

Furthermore, Supplementary Figures 5 and 6 show the DE analysis results with respect to different cell networks when we fixed the observed DE evidence and gene networks as the same in the Main models. With different cell network structures, the MRF models yielded comparable parameter estimates (Supplementary Figure 5). The UpSet plot in Supplementary Figure 6 shows the overlap of DEGs identified with three cell network structures. We note that the DE results were consistent across different cell networks with slight variations. In addition, the IPA enrichment results also demonstrate that our MRF models have fairly robust performance with respect to different cell network structures.

## Conclusions and Discussions

By borrowing information through known biological networks, our proposed method, MRFscRNAseq, provides differential expression analysis to identify celltype specific DEGs for scRNA-seq data sets with increased statistical power. With observed DE evidence, it utilizes a Markov Random Field model to appropriately take gene network information as well as dependencies among cell types into account. We implemented an Expectation-Maximization (EM) algorithm with mean field-like approximation to estimate model parameters and a Gibbs sampler to infer DE status. Simulation study showed that our method has better power to detect cell-type specific DEGs than conventional methods while appropriately controlling type I error rate. In the differential expression analysis using an IPF scRNA-seq data set, we have showed that our method is able to identify an additional set of novel DEGs using summary statistics from various existing differential expression methods. Pathway analysis with IPA also suggests that the additional set of pathways have biological meanings that are strongly correlated with IPF.

For gene networks, we utilized two protein-protein interaction network databases in the IPF scRNA-seq data application, BioGrid and IntAct. In fact, our method can be adapted to other networks that have similar structures as BioGrid or IntAct PPIN, i.e., KEGG pathways [66]. Furthermore, our model can be readily extended to many other existing DE methods with ease, just like what we have done with Wilcoxon test or MAST in the IPF scRNA-seq data analysis.

One caveat in our model is that the direction of changes in gene expressions is not directly incorporated in the model, which means that we are unable to differentiate whether these identified DEGs are up-regulated or down-regulated. One possible remedy is to use the sign of the original input test statistics to determine the sign of the DE results. For future work, weights could be added in our graphical model.

For instance, transcription factors probably should have more weights because of their importance in gene regulation.

## Supplementary Information

## Abbreviations

DE: Differential expression;
DEGs: Differentially expressed genes;
MRF: Markov Random Field;
EM: Expectation-Maximization;
IPF: Idiopathic pulmonary fibrosis;
scRNA-seq: single-cell RNA-sequencing.
KEGG: Kyoto Encyclopedia of Genes and Genomes;
GWAS: Genome-wide association studies;
PPIN: Protein-protein interaction network;
MAST: Model-based Analysis of Single-cell Transcriptomics.

## Authors’ contributions

HL and HZ designed the method and wrote the manuscript. HL implemented the algorithm and wrote the R package. BZ helped with the simulation analysis. ZX helped with the IPF scRNA-seq analysis. TA and NK provided the IPF scRNA-seq data set and revised the manuscript. All authors read and approved the final manuscript.

## Funding

Supported in part by NIH grant GM134005 and NSF grant DMS 1902903 (HZ). NIH NHLBI grants R01HL127349, R01HL141852, U01HL145567, UH2HL123886 to NK, and a generous gift from Three Lakes Partners to NK.

## Availability of data and materials

The IPF scRNA-seq data set is available in the Gene Expression Omnibus repository (GSE136831) (https://www.ncbi.nlm.nih.gov/geo/query/acc.cgi?acc=GSE136831). The R package MRFscRNAseq and supplementary results is available on GitHub (https://github.com/eddiehli/MRFscRNAseq).

## Ethics approval and consent to participate

Not applicable.

## Consent for publication

Not applicable.

## Competing interests

NK served as a consultant to Biogen Idec, Boehringer Ingelheim, Third Rock, Pliant, Samumed, NuMedii, Theravance, LifeMax, Three Lake Partners, Optikira, Astra Zeneca, Veracyte, Augmanity and CSL Behring, over the last 3 years, reports Equity in Pliant and a grant from Veracyte, Boehringer Ingelheim, BMS and non-financial support from MiRagen and Astra Zeneca. NK has IP on novel biomarkers and therapeutics in IPF licensed to Biotech. The authors declare that they have no other competing interests.

## Acknowledgements

Not applicable.

## Notes

### Competing Interest Statement

The authors have declared no competing interest.

### Summary of Updates

We revised the simulation study section and data analysis section significantly.

